# Transcriptional analysis of sodium valproate in a serotonergic cell line reveals gene regulation through both HDAC inhibition-dependent and independent mechanisms

**DOI:** 10.1101/837732

**Authors:** Priyanka Sinha, Simone Cree, Allison L. Miller, John F. Pearson, Martin A. Kennedy

## Abstract

Sodium valproate (VPA) is a histone deacetylase (HDAC) inhibitor, widely prescribed in the treatment of bipolar disorder, and yet the precise modes of therapeutic action for this drug are not fully understood. After exposure of the rat serotonergic cell line RN46A to VPA, RNA-sequencing (RNA-Seq) analysis showed widespread changes in gene expression. Analysis by multiple pipelines revealed as many as 230 genes were significantly upregulated and 72 genes were significantly downregulated. A subset of 23 differentially expressed genes was selected for validation using the nCounter^®^ platform, and of these we obtained robust validation for *ADAM23, LSP1, MAOB, MMP13, PAK3, SERPINB2, SNAP91, WNT6*, and *ZCCHC12*. We investigated the effect of lithium on this subset and found four genes, *CDKN1C, LSP1, SERPINB2* and *WNT6* co-regulated by lithium and VPA. We also explored the effects of other HDAC inhibitors and the VPA analogue valpromide on the subset of 23 selected genes. Expression of eight of these genes, *CDKN1C, MAOB, MMP13, NGFR, SHANK3, VGF, WNT6* and *ZCCHC12*, was modified by HDAC inhibition, whereas others did not appear to respond to several HDAC inhibitors tested. These results suggest VPA may regulate genes through both HDAC-dependent and independent mechanisms. Understanding the broader gene regulatory effects of VPA in this serotonergic cell model should provide insights into how this drug works and whether other HDACi compounds may have similar gene regulatory effects, as well as highlighting molecular processes that may underlie regulation of mood.

## Introduction

Sodium valproate (VPA) is a histone deacetylase inhibitor (HDACi) [1, 2], and is widely used in the treatment of bipolar disorder (BD) [3]. The mechanism of action of VPA as a mood stabilizer is not well understood. VPA has been shown to affect the serotonergic [4, 5], dopaminergic [6], GABAergic [7] and glutamatergic [8–10] pathways as well as intracellular signalling pathways such as those mediated by phosphoinositol [11, 12], glycogen kinase-3β/Wnt [13], protein kinase C [14], and ERK/MAPK [15, 16] among others [17–19].

Previous attempts to find differential gene expression effects of the drug have focused mostly on a handful of candidate genes [6, 20–25]. Subsequently, microarrays were used to study the gene expression effects of VPA in neuroblastoma or glioma cell lines [26–32]. Microarrays are limited to identifying transcripts targeted by their probe sets, limiting their ability to detect novel transcripts. In contrast, RNA-Seq has the capacity to capture transcripts from both protein-coding and non-coding genes, which have been shown to be important transcriptional regulators. VPA has been shown to target multiple genes and pathways [26, 27, 31, 33-–36], therefore a whole transcriptomic approach is well suited to better understand potential mechanisms of action.

Dysregulation of the serotonergic system has been implicated in the pathophysiology of mood disorders [18]. The serotonergic neurons originate from the median and the dorsal raphe nucleus in the brain stem and project to different brain regions, where they secrete serotonin which regulates mood [37]. In this study, we examined the gene expression effects of VPA using the neural cell line RN46A, derived from the rat medullary raphe region [38], which has been used previously to study gene expression in response to antidepressants and mood stabilisers [39–44]. Differentiated RN46A cells express all 5-HT receptors 5-HT1A, 5-HT1B, 5-HT2A and 5-HT2C and the serotonin transporter SERT [38, 45].

Understanding the transcriptional effects of VPA in a relevant cellular context may provide clues to its mechanisms of action. Previous studies in our laboratory using the RN46A cell line revealed differential gene expression in response to VPA exposure [41, 44]. In this paper, we sought to extend these findings using RNA-Seq as a discovery tool for differentially expressed genes (DEG), followed by validation of a subset of these genes using an orthogonal gene expression analysis method. Furthermore, we examined the action of other mood stabilizing drugs and other HDACi compounds on expression of this subset of genes.

## Methods

### Cell culture and RNA extraction

The RN46A cells were maintained at 33°C in Dulbecco’s Modified Eagle Medium: Nutrient Mixture F-12 (DMEM/F12; ThermoFisher Scientific, MA, USA) containing glutamine (GlutaMAX™-I; ThermoFisher Scientific, MA, USA) and supplemented with 5% fetal bovine serum (FBS) and 250 μg/ml Geneticin G418 (ThermoFisher Scientific, MA, USA). For differentiation, RN46A cells were seeded in plates coated with 100μg/cm^2^ collagen (Sigma-Aldrich, St. Louis, MO, USA) and 1μg/cm^2^ fibronectin (Sigma-Aldrich, St. Louis, MO, USA), and confluent RN46A cultures were shifted to 39°C and supplemented with DMEM/F-12 containing 1% FBS, 1% BSA, 1% N2 supplement, 0.75 g/L sodium pyruvate and 0.073 g/L L-glutamine. RN46A cells were differentiated for 72 h and then drug exposed for 72 h.

Differentiated RN46A cells were exposed to either 0.5mM VPA or 0.5mM lithium chloride for 72 h. Three factors guided the concentration of lithium and VPA chosen in this study: plasma levels measured in patients undergoing treatment [8, 46–48], prior levels used in published *in vitro* [49–53] and *in vivo* studies [54–57], and prior levels used in studies from our laboratory [41, 44]. For RNASeq, cells cultured with differentiation medium were used as control.

Total RNA was isolated using Trizol^®^ (ThermoFisher Scientific, MA, USA), according to the manufacturer’s instructions. Quality control was performed using the Agilent 2200 TapeStation system (Agilent Technologies, Santa Clara, CA, USA). RNA samples with 260/280 and 260/230 ratios of ~2 and RIN score greater than 7 were shipped to Novogene (Beijing, China) for RNA-Seq. The drug exposure experiments were repeated three times and RNA samples from all three experiments were sent for sequencing.

For validation of RNA-Seq data, the nCounter^®^ platform (NanoString, Seattle, WA, USA) was used in two successive runs, each using RNA derived from independent cell culture experiments. VPA is a major class I HDACi [1] so in the first run, we tested whether a similar broad range HDACi (trichostatin A) could affect gene expression, and in the second run, selective HDAC 1, 3 and 8 inhibitors were used. In Run 1, differentiated RN46A cells were exposed to 0.5mM VPA, 1mM lithium, 0.5mM valpromide (VPD) and 30nM trichostatin A (TSA) (Supplementary Table 1). In Run 2, differentiated RN46A cells were exposed to 0.5mM VPA, 2mM lithium, 0.5mM VPD, 0.5μM CI994 (HDAC1 inhibitor), 0.08μM RGFP966 (HDAC3 inhibitor),, 0.01μM PCI-34051 (HDAC8 inhibitor) and 0.015μM tubastatin A (HDAC6 inhibitor) for 72 h (Supplementary Table 1). For all nCounter^®^ experiments, cells cultured with differentiation medium containing 0.05% DMSO were used as controls.

For nCounter^®^ experiments, direct-zol™ RNA miniprep kit (Zymogen, Irvine, CA, USA) was used for total RNA extraction according to the manufacturer’s instructions. nCounter^®^ mRNA gene expression assays were carried out by New Zealand Genomics Limited (Dunedin, New Zealand). Based on the nCounter^®^ custom code design, the drug exposure experiments were repeated four times for Run1 samples and three times for Run2 samples.

### RNA-Seq analysis

RNA-Seq was carried out on the Illumina HiSeq™ 2500 platform (Illumina inc. San Diego, CA, USA) by Novogene (Beijing, China). 125 bp paired-end sequencing generated >16 million reads per sample, which were mapped to the Rat Rnor6 reference genome (ENSEMBL, Jul 2014, version 88) with the splice-aware STAR aligner (v2.5.1.b) [58]. In addition, reads were mapped to the rat transcriptome with kallisto (v0.42.4) [59] and salmon (v0.6.1) [60]. Mapping quality data using STAR, kallisto and salmon software are summarized in Supplementary Table 2, 3 and 4, respectively.

Four pipelines were used for DEG analysis (Figure 1): DESeq2 [61], cufflinks2.2.1 suite [62, 63], kallisto-sleuth [59] and salmon-sleuth [59, 60]. Salmon output was converted into sleuth-compatible format using wasabi [60]. Genes with a cut-off adjusted p-values (padj) < 0.05 and log2 fold change (Log2FC) ≥ 1.5 fold were considered to be differentially expressed.

**Figure 1.**
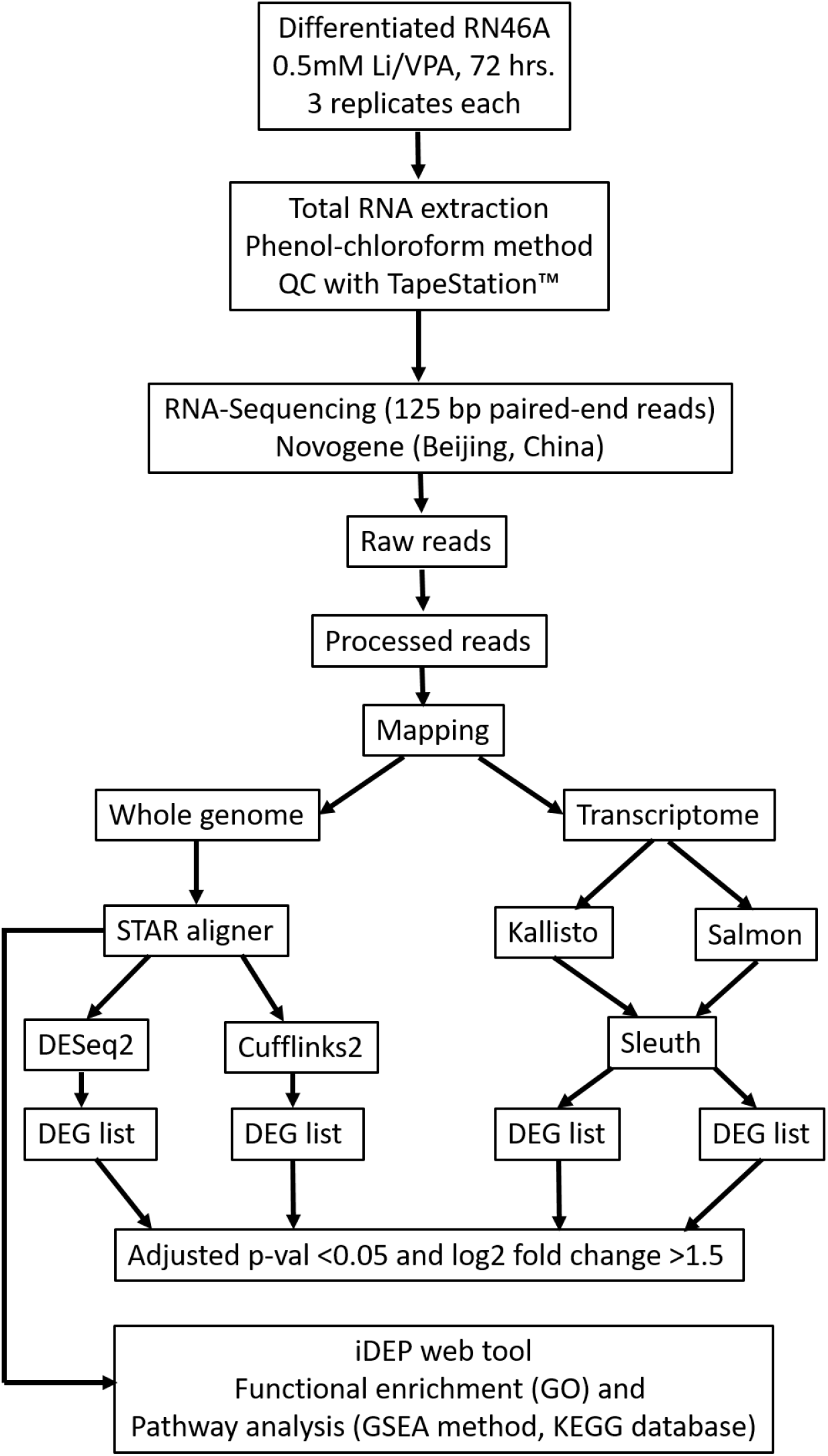
Flowchart of RNA-Seq data analysis procedure. Differential gene expression analysis was performed with DESeq2, cufflinks2 suite and sleuth. Li: lithium, QC: quality control, DEG: differentially expressed genes, iDEP: integrated differential expression and pathway analysis. GSEA: gene set enrichment analysis, KEGG: Kyoto Encyclopedia of Genes and Genomes.

### Functional enrichment and pathway analysis

Integrated differential expression and pathway analysis (iDEP) web-based tool (v 0.80) [64] was used for functional enrichment and pathway analysis. Read count data from STAR aligner for three VPA-treated and three untreated control samples were uploaded to the website. Differential expression analysis was carried out with DESeq2. The count data were rlog transformed [61] using the edgeR [65] tool for further clustering analysis and principal component analysis. Enrichment analysis was carried out on DEG from the VPA exposure experiments using GO and available gene sets with the iDEP web tool [64]. Pathway analysis was carried out with the GSEA method and KEGG database using the iDEP web tool.

### nCounter^®^ data analysis

Raw data from the reporter code count (RCC) files were analysed with nSolver^TM^ v3.0. The mRNA counts were normalized with a geometric mean of positive controls, geometric mean of negative controls and geometric mean of housekeeping genes (*G6PD* and *MAPK6*). One-tailed heteroscedastic Welch’s t-test was used for DE analysis.

The R package, NanoStringDiff (v 1.8.1), was used for normalization of raw nCounter^^®^^ data and DE analysis [66]. Empirical Bayes shrinkage was used to estimate dispersion parameters and counts were modeled using the negative binomial distribution. A generalized linear model (GLM) maximum likelihood ratio test was used for DE analysis and chi-square approximation for calculating p-values. The Benjamini-Hochberg method was used for calculating FDR [67].

## Results

### Differential gene expression analysis in response to VPA and lithium

In this study, RNA-Seq was used to examine global gene expression changes in differentiated RN46A cells exposed to VPA and lithium. A total of 19,774 genes from STAR alignment were used for differential expression analysis with DESeq2 and cufflinks2 suite whereas ~10,394 and ~10,134 genes were used after pseudo-alignment with kallisto-sleuth and salmon-sleuth, respectively. Genes with padj < 0.05 and Log2FC ≥ 1.5 fold were considered as significantly regulated (DEG). Using these criteria, in response to VPA, 224 DEGs were observed with DESeq2 (Supplementary Table 5), 144 with kallisto (Supplementary Table 6), 158 with salmon (Supplementary Table 7) and 302 with cuffdiff2 (Supplementary Table 8). A four-way comparison showed that a total of 67 DEGs were common among the four tools, whereas 151 genes were common between the whole genome (DESeq2 and cuffidiff2) and 111 genes between the transcriptome (salmon and kallisto) differential expression analysis tools (Figure 2).

**Figure 2.**
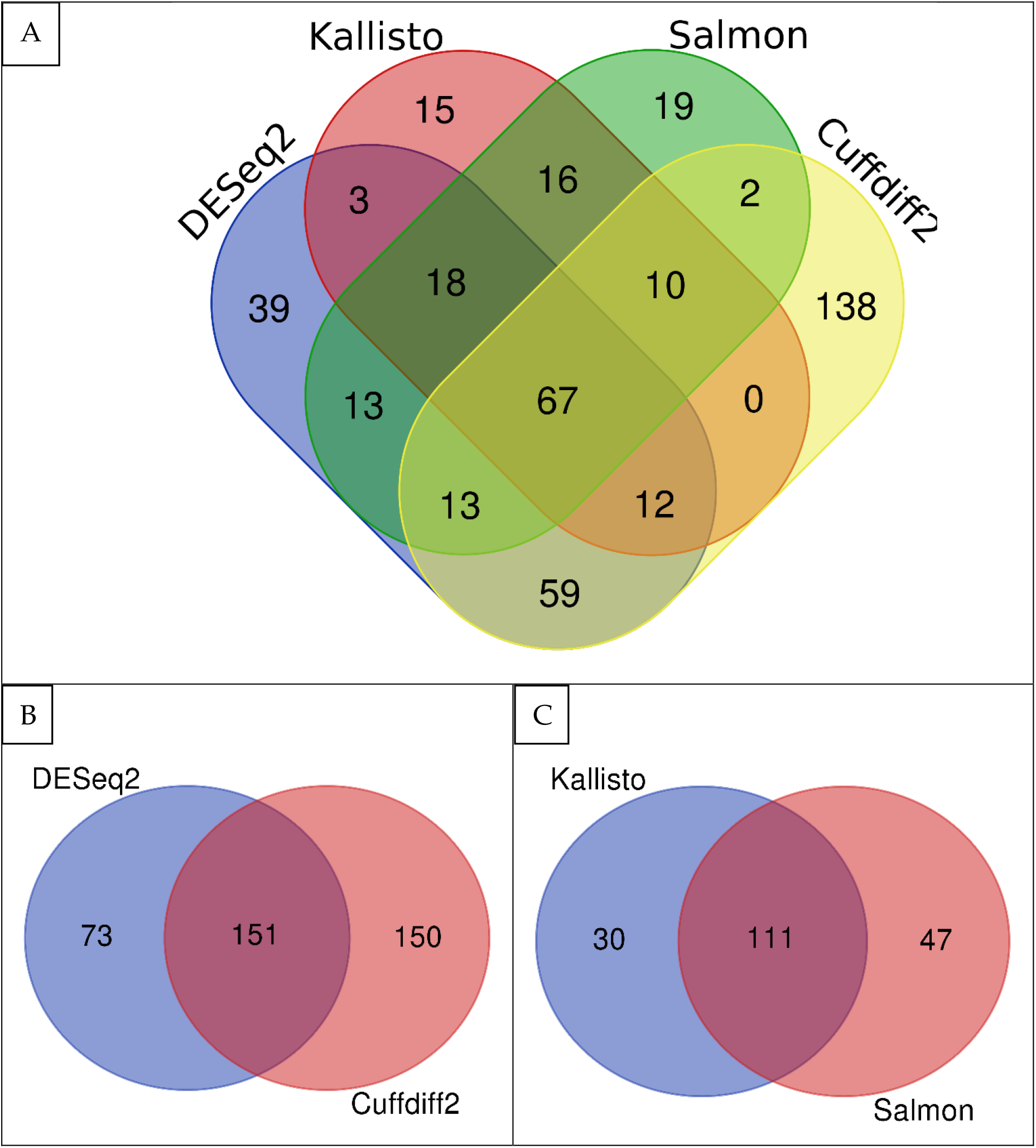
Agreement between the four DEA methods. The panel A shows the four-way comparison of the DEA tools. The panels B and C indicate the two-way comparison to clarify relationships between whole-genome DEA methods (Cuffdiff2 and DESeq2) and transcriptome DEA methods (kallisto and salmon).

The top 25 DEGs in response to 0.5 mM VPA as identified with DESeq2 are listed in Table 1. In comparison, exposure of RN46A cells to 0.5 mM lithium resulted in no genes showing significant expression differences in the RNA-Seq analysis.

**Table 1.** Top 25 differentially expressed genes in response to VPA as identified with DESeq2.

| Gene name | Base mean control<sup>1</sup> | Base mean VPA<sup>1</sup> | log2FC | padj<sup>2</sup> |
| --- | --- | --- | --- | --- |
| *IGF2* | 1.78 ± 0.71 | 139.77 ± 17.44 | 4.15 | 8.40E-32 |
| *ADGRB2* | 34.57 ± 11.33 | 418.11 ± 47.94 | 3.03 | 4.38E-28 |
| *AQP1* | 1813.82 ± 161.36 | 349.61 ± 34.24 | -2.26 | 7.43E-28 |
| *MMP13* | 1066.38 ± 144.15 | 6415.40 ± 855.59 | 2.39 | 1.22E-25 |
| *VGF* | 30.11 ± 17.49 | 642.32 ± 161.08 | 3.34 | 1.07E-24 |
| *ZCCHC12* | 60.32 ± 6.42 | 443.85 ± 73.79 | 2.52 | 1.24E-22 |
| *IL20RB* | 190.62 ± 35.73 | 14.04 ± 5.16 | -2.95 | 3.27E-20 |
| *NGFR* | 47.83 ± 8.42 | 327.14 ± 37.71 | 2.38 | 1.79E-19 |
| *TSPAN13* | 72.98 ± 8.75 | 457.96 ± 87.42 | 2.33 | 1.86E-18 |
| *COL3A1* | 2014.23 ± 727.89 | 245.37 ± 50.45 | -2.53 | 4.17E-17 |
| *LSP1* | 186.31 ± 12.93 | 22.84 ± 8.96 | -2.61 | 4.63E-17 |
| *AGAP2* | 0.25 ± 0.44 | 63.11 ± 9.77 | 3.31 | 1.34E-16 |
| *SNAP91* | 13.02 ± 3.28 | 137.50 ± 10.69 | 2.63 | 2.18E-16 |
| *WNT6* | 30.12 ± 6.36 | 227.07 ± 38.69 | 2.38 | 8.01E-16 |
| *NXPH4* | 3.74 ± 2.32 | 79.60 ± 18.76 | 2.98 | 8.42E-16 |
| *TMEM130* | 19.20 ± 5.1 | 162.75 ± 26.61 | 2.47 | 1.55E-14 |
| *ADAM23* | 19.14 ± 8.71 | 123.74 ± 24.97 | 2.55 | 1.91E-14 |
| *CLGN* | 30.93 ± 2.57 | 176.08 ± 24.33 | 2.20 | 8.31E-14 |
| *ERBB3* | 67.54 ± 16.86 | 289.04 ± 22.42 | 1.99 | 3.36E-13 |
| *RGD1563349* | 5.10 ± 1.49 | 67.82 ± 5.15 | 2.73 | 7.39E-13 |
| *CHRNA5* | 2.48 ± 1.86 | 52.68 ± 6.89 | 2.86 | 1.30E-12 |
| *ATP1B2* | 38.83 ± 8.26 | 199.63 ± 29.11 | 2.11 | 1.34E-12 |
| *TMEFF1* | 1.88 ± 1.46 | 53.25 ± 5.51 | 2.87 | 1.74E-12 |
| *FAM25A* | 82.84 ± 33.0 | 5.53 ± 1.74 | -2.71 | 2.06E-12 |
| *PERP* | 26.70 ± 4.62 | 150.22 ± 23.25 | 2.16 | 3.29E-12 |

<sup>1</sup>Ranked by p-values. Base mean control and base mean VPA refer to the mean of normalized counts from four replicates each of untreated and VPA-treated samples. Fold difference between the untreated (control) samples and samples treated with VPA are shown.
<sup>2</sup>FDR indicates false discovery rates which are essentially p values adjusted for multiple testing using the Benjamini-Hochberg procedure. padj- adjusted p-value.

### Functional enrichment and pathway analysis

iDEP web-based tool v0.80 [64] was used for functional enrichment and pathway analysis for VPA exposure. Read count data from STAR-aligner was analyzed with DESeq2 and a total of 985 genes was selected at FDR of 0.1 and fold change of 2, with 712 up-regulated and 273 down-regulated genes. The most significant GO processes and pathways are summarized in Supplementary Table 9 for upregulated genes and Supplementary Table 10 for downregulated genes. Both the GO processes and pathways for the upregulated genes were highly enriched in neuronal development and function. For the downregulated genes, the GO processes were enriched in response to environmental stimuli, however, the pathways were enriched in chromatin and nucleosome assembly (associated with HDACi effects of VPA).

### Validation of RNA-Seq results with nCounter^®^ assay

The nCounter^®^ assay format chosen for independent validation of DEGs identified in RNA-Seq required that we focus on a relatively small subset of DEG. Therefore, 23 VPA-regulated genes (Supplementary Table 11) were selected for validation. The selection process began with those genes that showed high fold change and strong statistical significance in the RNA-Seq data. The expression characteristics of each candidate DEG for validation were then examined in the Genotype-Tissue Expression (GTEx) [68] and Allen brain atlas *in-situ* hybridisation [69] databases, and brain-expressed genes were preferentially selected (Supplementary Table 12).

Of the 23 genes in the nCounter^®^ assay panel, 12 genes were found to be altered by VPA in Run1 and 17 genes in Run2 as identified with both analysis methods (Table 2). Taken together, six genes replicated as VPA DEGs in both Run1 and Run2, using both nSolver™ and nanoStringDiff software (Table 2). These were *SNAP91, ADAM23, LSP1, ZCCHC12, PAK3* and *MMP13*. Other genes with high confidence included *MAOB, SERPINB2* and *WNT6*. All the gene expression changes were in the same direction between RNA-Seq and nCounter^®^ platforms, and showed similar magnitude.

**Table 2.**
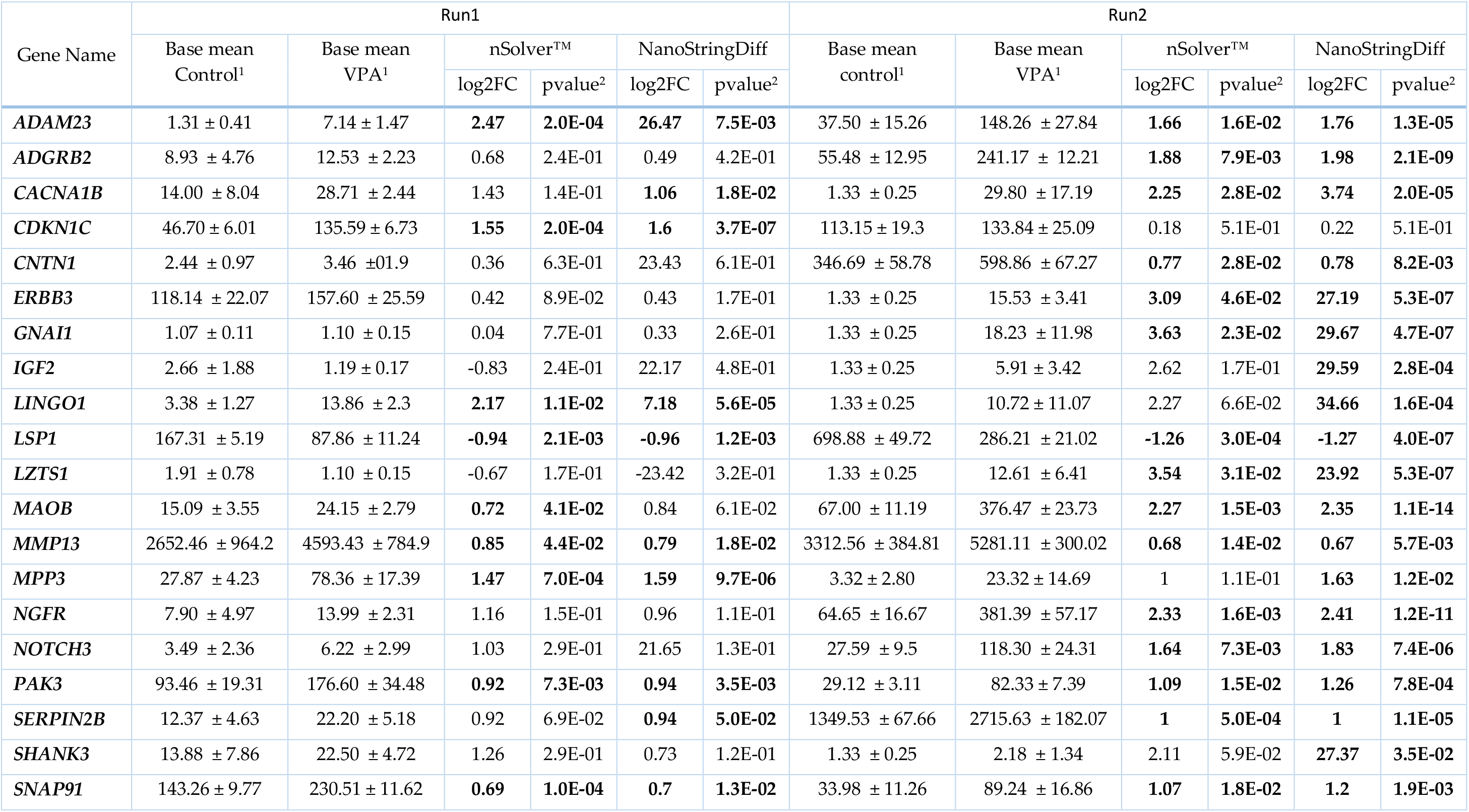
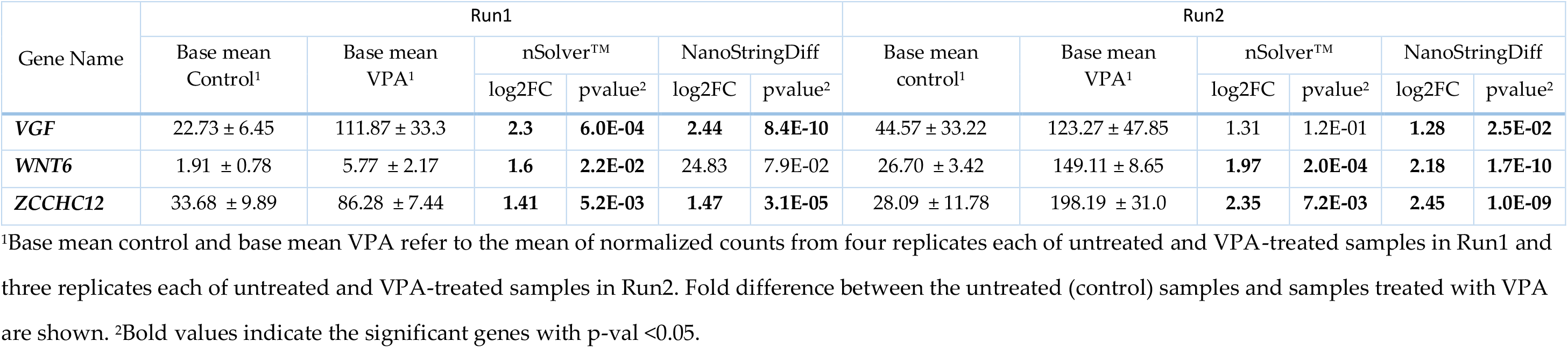
Genes showing significant expression difference after VPA exposure (Run1 and Run2)

### Lithium

No significant gene expression changes in the RNA-Seq data were observed with exposure of the RN46A cell line to 0.5mM lithium. This concentration of lithium was judged to be quite low for RN46A cells, and therefore was increased to 1mM and 2mM in nCounter^®^ Run1 and Run2, respectively. After exposure to 1mM lithium in Run 1, expression of *CDKN1C* was significantly changed and in Run 2, 2mM lithium resulted in significant expression changes for three genes *(LSP1, SERPINB2* and *WNT6*) (Table 3). When analysed with NanoStringDiff, two genes (*CDKN1C* and *SERPIN2B*) showed altered expression in Run1 and six genes (*LSP1, SERPIN2B, WNT6, MAOB, CNTN1* and *ADAM23*) showed altered expression in Run2.

**Table 3.** Genes showing significant expression difference after lithium exposure (Run1 and Run2).

| Gene Name | Run1 (1.0 mM lithium) |  |  |  |  |  | Run2 (2.0 mM lithium) |  |  |  |  |  |
| --- | --- | --- | --- | --- | --- | --- | --- | --- | --- | --- | --- | --- |
|  | Base mean control<sup>1</sup> | Base mean lithium<sup>1</sup> | nSolver™ |  | NanoStringDiff |  | Base mean control<sup>1</sup> | Base mean lithium<sup>1</sup> | nSolver™ |  | NanoStringDiff |  |
|  |  |  | log2FC | pvalue<sup>2</sup> | log2FC | pvalue<sup>2</sup> |  |  | log2FC | pvalue<sup>2</sup> | log2FC | pvalue<sup>2</sup> |
| *CDKN1C* | 46.70 ± 6.01 | 75.97 ± 10.25 | **0.7** | **3.7E-03** | **0.71** | **5.47E-05** | 113.15 ± 19.3 | 84.00 ± 20.19 | -0.4 | 2.5E-01 | -0.4 | 1.8E-01 |
| *LSP1* | 167.31 ± 5.19 | 204.42 ± 19.01 | 0.28 | 2.9E-02 | 0.28 | 3.23E-03 | 698.88 ± 49.72 | 480.36 ± 16.73 | **-0.53** | **7.0E-03** | **-0.53** | **4.6E-02** |
| *SERPIN2B* | 12.37 ± 4.63 | 6.72 ± 0.76 | -0.77 | 1.0E-01 | **-1.85** | **8.78E-03** | 1349.53 ± 67.66 | 2489.91 ± 273.19 | **0.87** | **7.0E-03** | **0.88** | **9.1E-04** |
| *WNT6* | 1.91 ± 0.78 | 1.51 ± 0.43 | -0.26 | 5.8E-01 | -16.82 | 4.78E-01 | 26.70 ± 3.42 | 4.92 ± 5.37 | **-1.32** | **5.0E-02** | **-1.6** | **3.4E-04** |
| *MAOB* | 15.09 ± 3.55 | 17.81 ± 4.66 | 0.24 | 4.6E-01 | 0.19 | 6.28E-01 | 67.00 ± 11.19 | 106.67 ± 20.64 | 0.57 | 6.4E-02 | **0.61** | **4.2E-02** |
| *CNTN1* | 2.44 ± 0.97 | 5.09 ± 2.7 | 0.95 | 2.1E-01 | 21.75 | 4.36E-01 | 346.69 ± 58.78 | 235.92 ± 28.77 | -0.52 | 8.0E-02 | **-0.54** | **5.0E-02** |
| *ADAM23* | 1.31 ± 0.41 | 3.42 ± 3.58 | 0.77 | 3.9E-01 | -80.18 | 4.26E-01 | 37.50 ± 15.26 | 13.43 ± 4.78 | -0.86 | 8.9E-02 | **-1.1** | **3.2E-03** |

<sup>1</sup>Base mean control and base mean lithium refer to the mean of normalized counts from four replicates each of untreated and lithium-treated samples in Run1 and three replicates each of untreated and lithium-treated samples in Run2. Fold difference between the untreated (control) samples and samples treated with lithium are shown. <sup>2</sup>Bold values indicate the significant genes with p-val < 0.05.

### Gene regulation by HDAC inhibitors

The effects of several different HDACi on expression of the selected subset of genes were evaluated using the nCounter^®^ assay in two separate runs. Log2FC and p-value for the 23 genes after exposure of RN46A cells to VPA, VPD and TSA in Run 1 are shown in Table 4. Log2FC and p-value for the 23 genes after exposure of RN46A cells to VPA, VPD and CI994 in Run2 are shown in Table 5, and RGFP966, PCI34051 and tubastatin A in Run2 are shown in Supplementary Table 13. Genes with Log2FC > 0.5 and p-value <0.05 were considered to be differentially expressed (DEG).

**Table 4.**
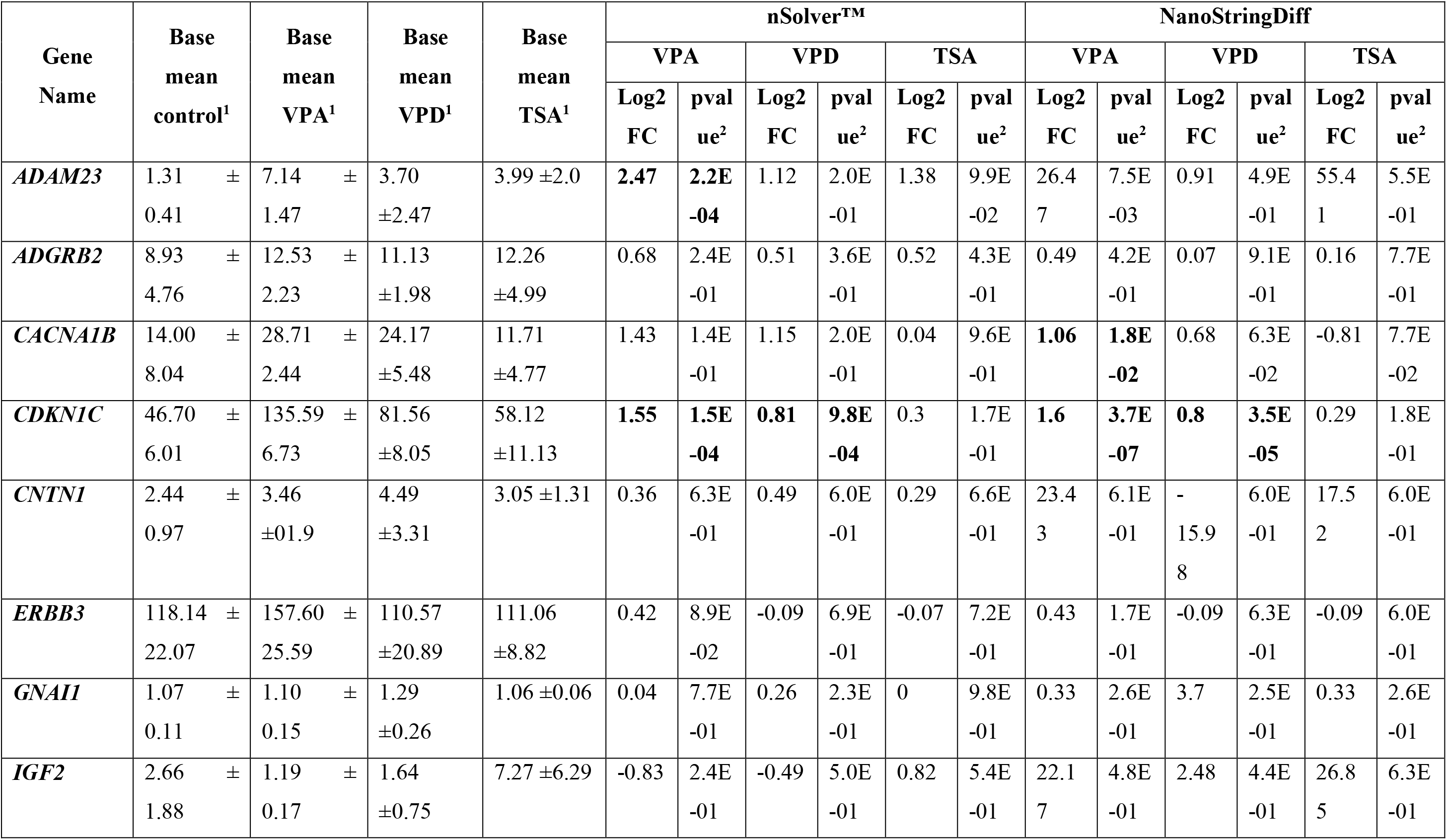
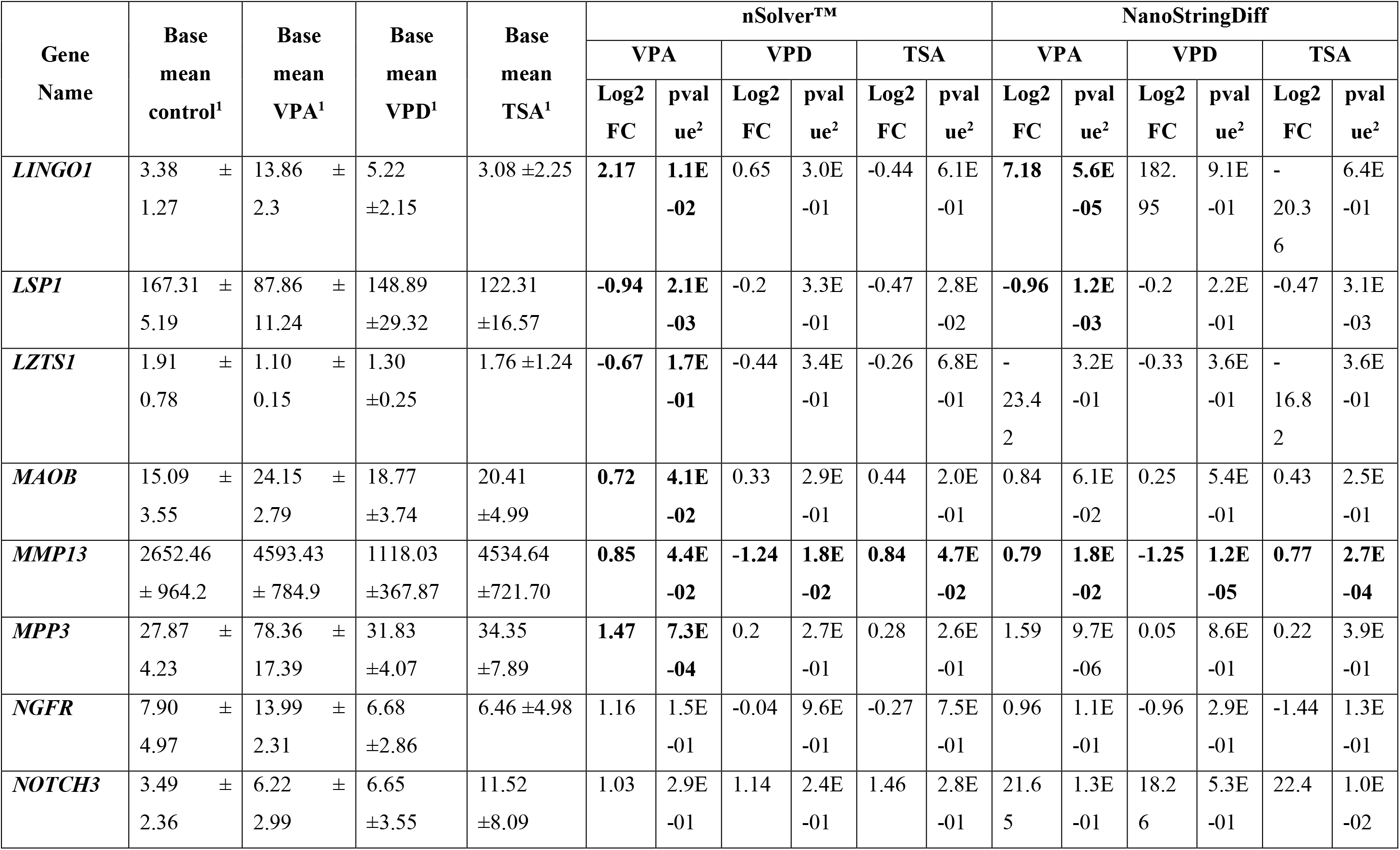
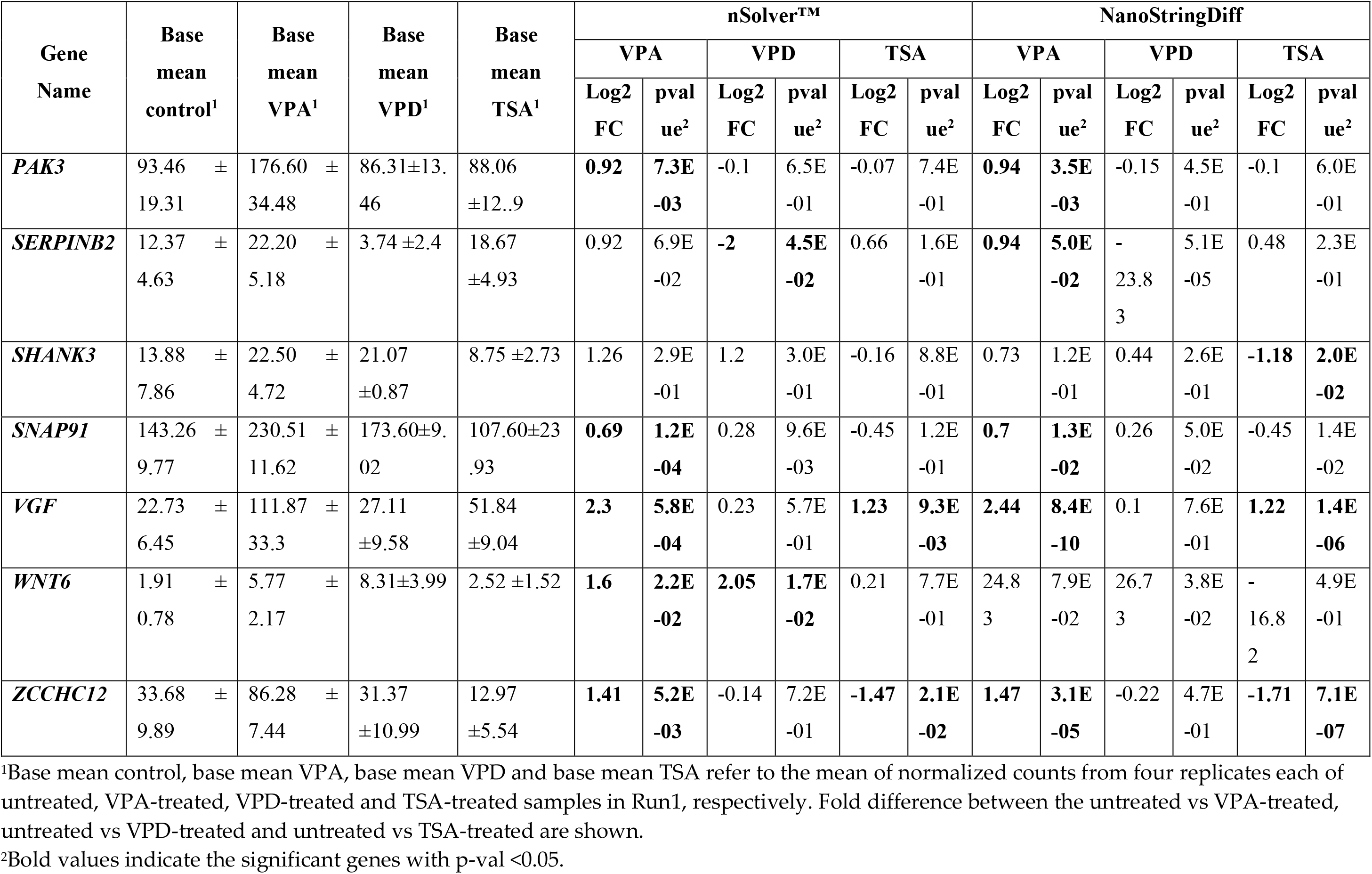
nCounter^®^ log2 fold change and p-values for VPA, VPD and TSA-treated RN46A cells in Run1.

**Table 5:**
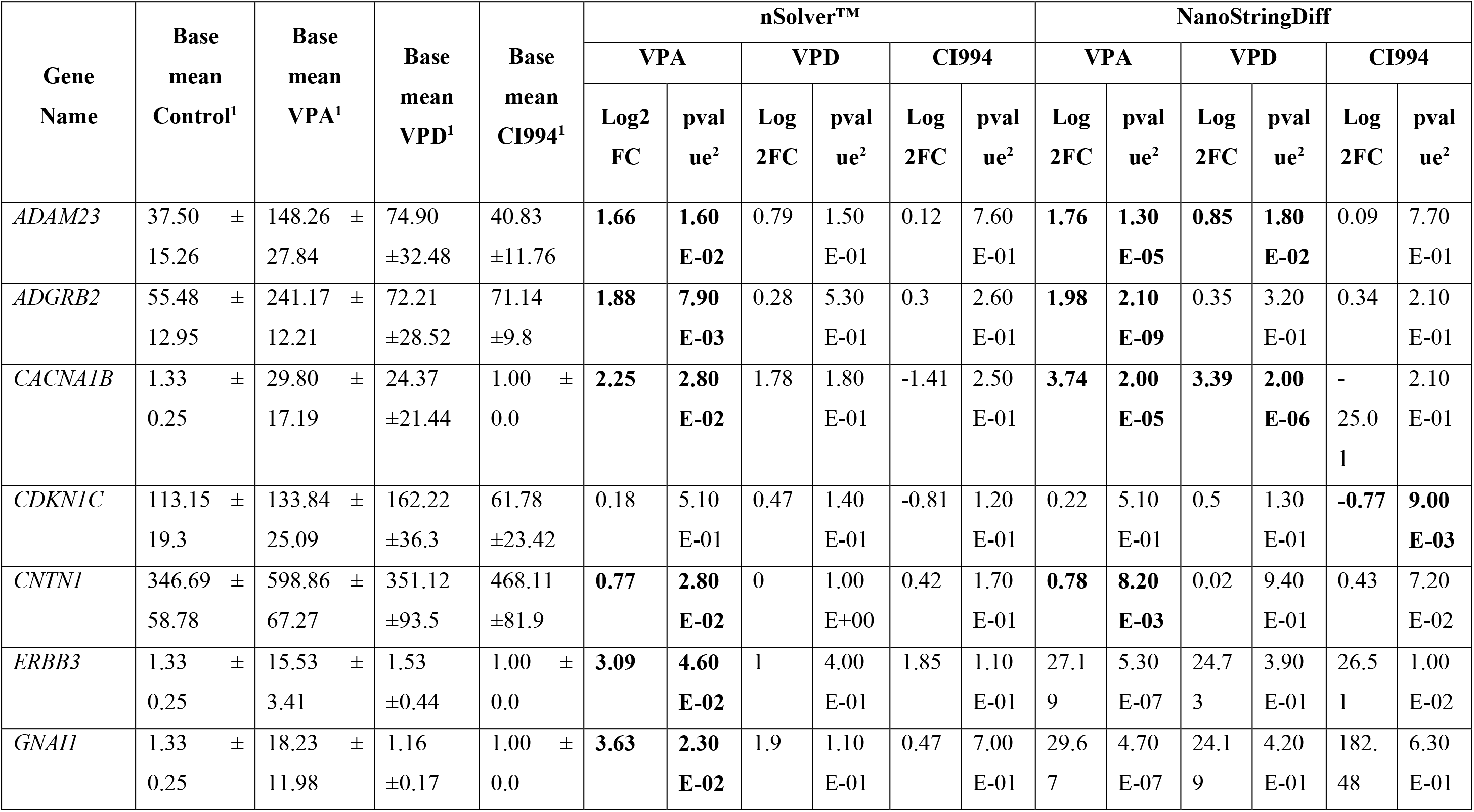
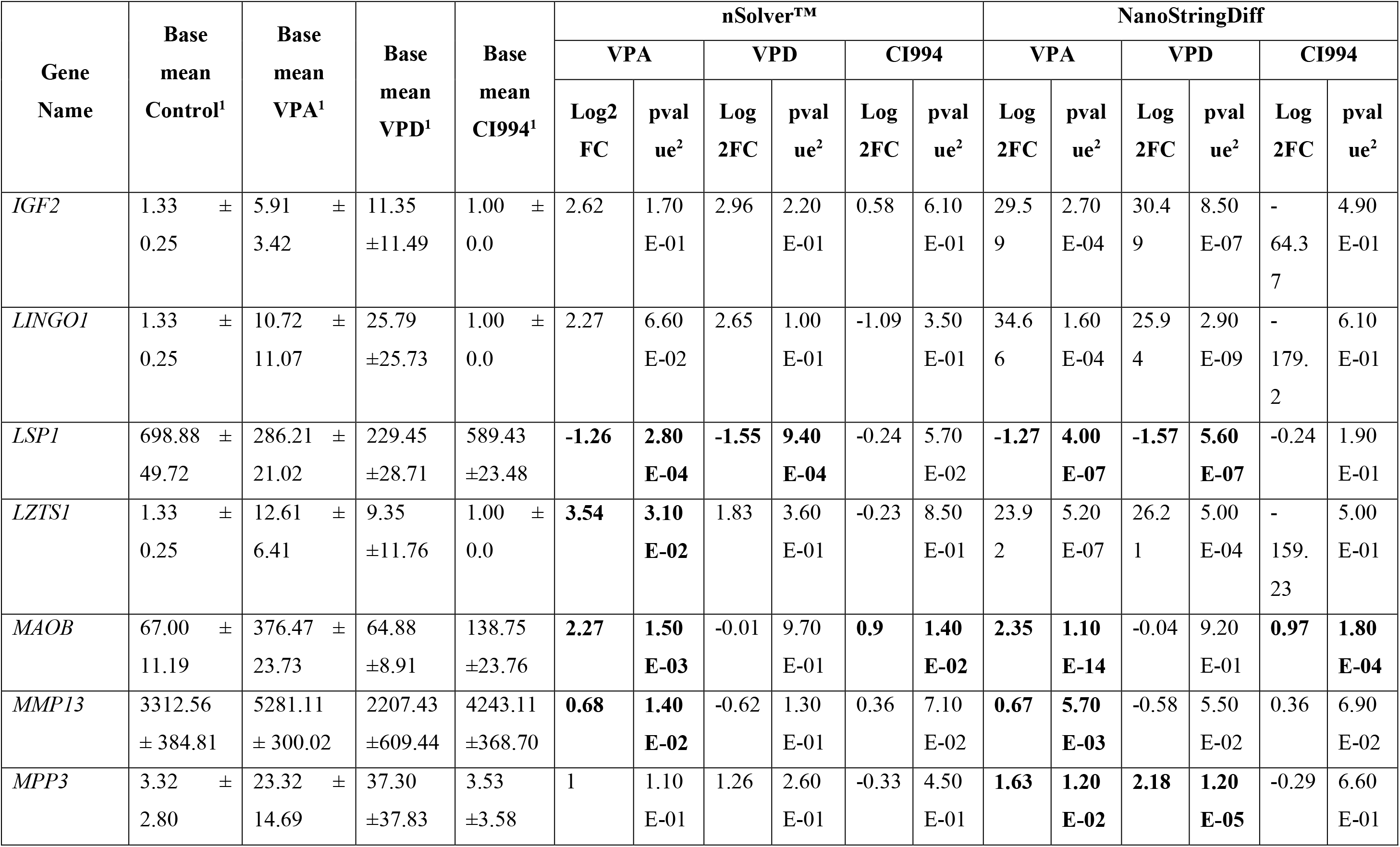
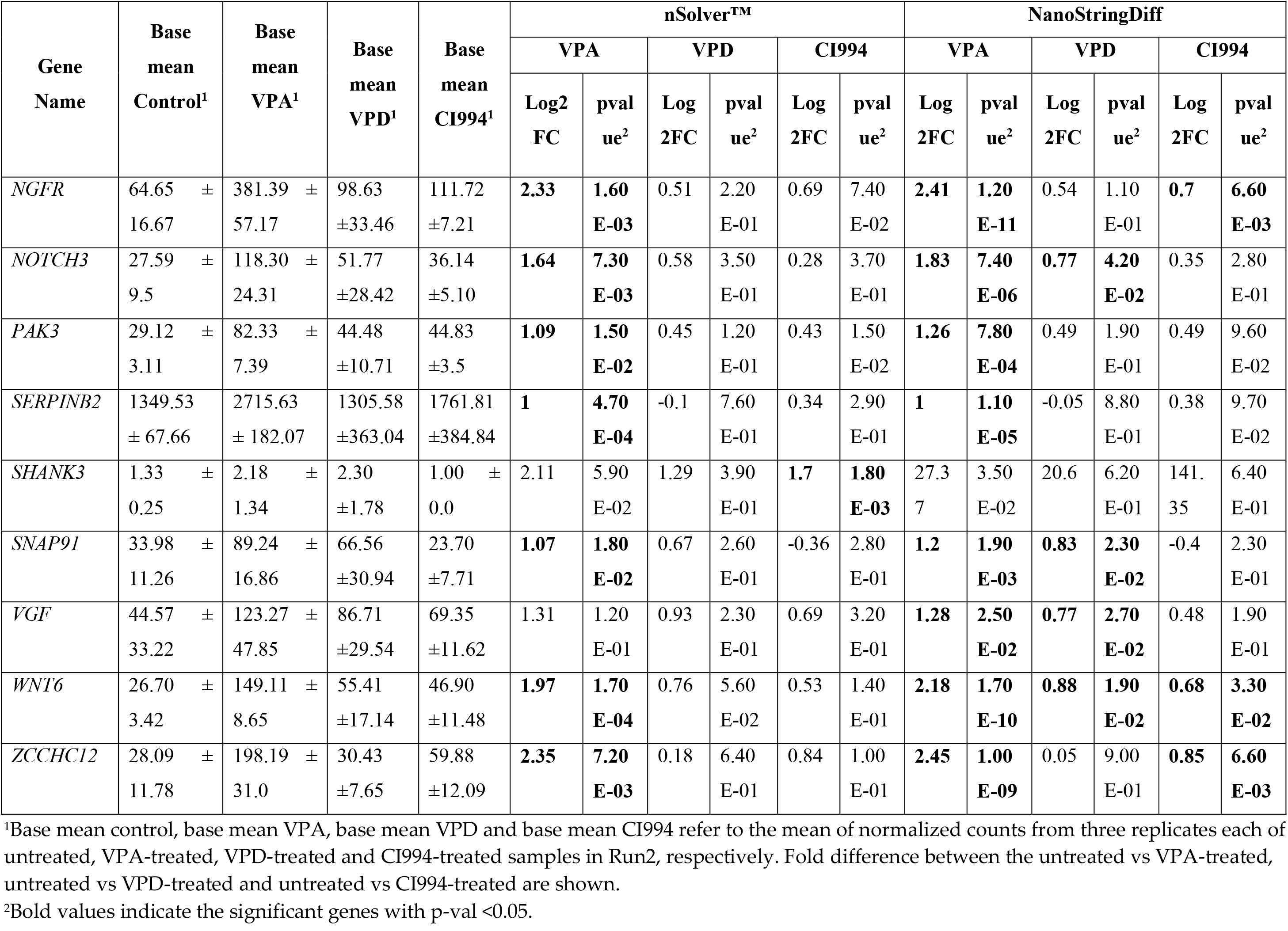
nCounter^®^ log2 fold change and p-values for VPA, VPD and CI994-treated RN46A cells in Run2.

After exposure to VPD, *CDKN1C* and *WNT6* were upregulated, and *MMP13* and *SERPINB2* were downregulated by VPD as identified with both nSolver^TM^ and NanoStringDiff in Run1 (Table 4). In Run2, only *LSP1* was a DEG as identified with both tools (Table 5). However, *ADAM23, CACNA1B, CDKN1C, IGF2, LINGO1, NOTCH3, SNAP91, VGF* and *WNT6* were identified as DEG with NanoStringDiff only.

*MMP13* and *VGF* were upregulated whereas *ZCCHC12* was downregulated in response to TSA as identified with both tools. *NOTCH3* was upregulated and *SHANK3* was downregulated by TSA as identified with NanoStringDiff only. After exposure to CI994, only *MAOB* was a DEG as identified with both nSolver^TM^ and NanoStringDiff (Table 5). However, *CDKN1C, NGFR*, *WNT6* and *ZCCHC12* were identified as DEG with NanoStringDiff only.

No DEG were identified for PCI34051 and tubastatin A exposure using either or both analysis tools (Supplementary Table 13). However, for RGFP966 exposure, *SNAP91* was identified as differentially expressed with NanoStringDiff only (Supplementary Table 13).

## Discussion

This study examined the global profile of gene expression effects of VPA and lithium, using RNA-Seq analysis, and more targeted effects of various drugs and HDACi on a subset of genes, in a serotonergic context. The RN46A cell line is derived from the medullary raphe nucleus, which along with the dorsal raphe nucleus is the serotonergic centre of the brain, which has been implicated in mood disorders and their treatment [70]. Upon differentiation, RN46A cells acquire a neuronally restricted morphology and express p75^NGFR^ and trkB receptors, neurofilament (NF-L, NF-M and NF-H), vimentin and nestin [38]. Therefore, we used this cell line as an *in vitro* model to explore expression changes that may occur within cells of the serotonergic system of the brain, after exposure to VPA.

The RNA-Seq analysis revealed that VPA resulted in extensive gene expression changes. To add confidence, we used four RNA-Seq analytical pipelines with freely-available and widely-used tools, and compared the different RNA-Seq workflows. There was variation in methodology among differential gene expression analysis tools including the choice of normalization method, density distribution and tests for differential expression such as t-test vs generalized linear models. Thus, each bioinformatic tool called different numbers of DEGs with cuffdiff2 producing the highest number, and kallisto and salmon producing similar numbers. A four-way comparison showed that a total of 67 DEGs were in common among the four DEA tools, and a two-way comparison to clarify relationships between the whole genome (and transcriptome) DEA tools showed that 151 DEGs were in common to DESeq2 and cuffdiff2, and 111 genes were in common to salmon and kallisto (Figure 2). The extensive gene expression changes seen with VPA in this model system are of interest. In part this may reflect the action of HDAC inhibitors on chromatin modifications, which has the potential to alter regulation of many genes. The widespread gene expression changes may also be relevant to the significant side effects that can occur with VPA, particularly teratogenicity [1, 71, 72].

In comparison, no significant gene expression changes were observed in the RNA-Seq data with lithium. A similar observation was made in a previous RNA-Seq analysis of lithium-treated undifferentiated RN46A cells [41]. With the same concentration and exposure time, only three genes were significantly altered with the same criteria (Log2FC >1.5 and FDR < 0.05). This suggested 0.5mM lithium may be too low to trigger gene expression changes in this cell line. In other cells, lithium concentrations as high as 10mM have been used [73], and in the SK-N-AS neuronal cell line, no change in synapsin 2a mRNA levels was seen at 0.5mM lithium, but it was differentially expressed with exposure to either 1.0 or 2.0 mM lithium [74].

Twenty-three DEGs from the RNA-Seq analysis of VPA-treated RN46A cells were selected for validation with the nCounter^®^ assay in two separate runs. Two software packages nSolver^TM^ v3.0 and nanoStringDiff were used for normalization and differential expression analysis. Twelve genes were found to be altered by VPA in Run1 and 17 genes in Run2 (Table 2) as identified with both analysis methods. Although there were some differences in DEG detected between the two runs, which were carried out independently, there was significant overlap and the direction of change was the same for all significant genes between runs, with comparable fold-changes. VPA results were reproducible across runs with slight variation due to the software used for analysis.

### Functional enrichment and pathway analysis

Functional enrichment analysis such as GO analysis characterizes a phenotype by identifying over-represented/enriched categories of genes that share a similar function within the list of DEGs. Biologically relevant signals may be detected by pathway analysis wherein genes are analysed in functional groups, networks or pathways.

The most-enriched GO processes for VPA regulated genes indicated the relevance to mood regulation of the genes identified in this RNA-Seq study (Supplementary Table 9). GO analysis highlighted biological processes “nervous system development” and “neurogenesis” as the most-enriched GO processes. Reduced neurogenesis has been linked to mood disorders [75, 76] and the therapeutic effect of antidepressants has been partly attributed to increased hippocampal neurogenesis [77–79]. Reduced hippocampal volume was observed in major depressive disorder and BD patients [80] which could be restored with mood stabilizers and antidepressants [81–83]. VPA has been shown to affect neuron differentiation and nervous system development [84–87], and similar biological processes such as neuron differentiation, projection and development were observed as the most-enriched in functional enrichment analyses of DEGs and differentially methylated regions in BD post-mortem brain [88].

VPA-regulated genes were analysed with the gene set enrichment analysis (GSEA) method [89] using the iDEP web tool. The top up-regulated pathway was cell adhesion molecules (Supplementary Table 9). ECM remodelling and synaptic plasticity dysfunction have been implicated in the pathophysiology of BD and schizophrenia [90–93]. Cell adhesion genes were most differentially regulated in response to VPA and matrix metalloproteinases such as *MMP13, MMP9* and *MMP12* were some of the genes highly upregulated by VPA. *MMP13* and *MMP12* were also upregulated by VPA in our prior RNA-Seq study with undifferentiated RN46A cells [41]. Other top up-regulated pathways were involved in GABAergic and serotonergic synapse suggesting that VPA up-regulated genes might be relevant to its mood-stabilizing effect.

### Co-regulated genes

Genes co-regulated by lithium and VPA, two structurally dissimilar mood stabilizers, are potentially relevant to mood stabilizing effects and may highlight important pathways in the mechanism of action for these drugs. Prior studies have identified some genes and pathways that overlap between lithium and VPA [32, 41, 94–96]. Four genes were clearly identified to be co-regulated by lithium and VPA: *CDKN1C*, *LSP1, SERPINB2* and *WNT6*. *CDKN1C* is a maternally imprinted gene regulated by HDAC inhibitors [97] and linked with neurogenesis in the developing brain [98]. *LSP1* is expressed in monocytes, neutrophils, macrophages as well as neuropils and may be linked to neuroinflammation in the brain. Both mania and depression are associated with increased pro-inflammatory cytokines secreted by macrophages, T-lymphocytes and endothelial cells, and decreased anti-inflammatory markers [99–103]. *SERPINB2* is involved in cell proliferation, differentiation and growth and regulates synaptic plasticity in hippocampal neurons [104]. Reduced synaptic plasticity is observed in BD [105, 106] and antidepressants have been shown to enhance neuroplasticity and resilience [77–79]. Wnt proteins regulate several processes such as brain development, cell growth, differentiation, migration and fate determination among others, implicated in BD pathophysiology. Both lithium and VPA affect expression of Wnt signalling pathway genes [107, 108]. Further study is needed to examine the role of these co-regulated genes in mood regulation.

### Gene regulation by other HDAC inhibitors

It is well-recognized that VPA has a range of pharmacological properties, including HDAC inhibition (HDACi). We examined whether other HDAC compounds have similar gene expression signatures as VPA and which specific HDAC isoform contributes to gene regulation. In the first nCounter^®^ experiments, VPA-regulated genes *MMP13, VGF* and *ZCCHC12* showed differential expression when exposed to TSA, thus implying HDAC inhibition might be important in regulation of these genes. VPA is a major class I HDACi [1] therefore, in the second nCounter^®^ run, selective HDAC 1, 3 and 8 inhibitors were used. Gene expression changes in response to VPA, VPD (a non-HDACi analogue of VPA) and other HDAC inhibitors were compared, and genes observed in both runs, with either or both analyses, were considered to be reproducible and of high confidence.

Comparing DEG between VPA and VPD exposures, *MMP13* and *SERPINB2* were downregulated by VPD but upregulated by VPA. For HDAC inhibitors, changes were only observed with TSA and CI994 (HDAC 1 inhibitor). TSA upregulated *MMP13* and *VGF*, and downregulated *ZCCHC12*. CI994 upregulated *MAOB, NGFR, WNT6* and *ZCCHC12*. Notably, the direction of change differed between these two drugs for *ZCCHC12*; VPA and CI994 upregulated *ZCCHC12* whereas TSA downregulated it.

This study revealed important insights into the mechanism of action of VPA. Comparing gene expression with VPA, other HDACs and a non-HDACi analogue of VPA shows that the pattern of DEGs regulated by VPA is probably a sum of its different mechanisms of action. VPA-regulated DEGs are only partly shared with another HDAC inhibitor or VPD. Our findings therefore suggest that VPA may have multiple mechanisms of action, with both its HDACi and other properties being important.

No change in gene expression was observed with HDAC3 (RGFP966) and HDAC8 inhibitors (PCI34051), which suggested that the HDAC3 and HDAC8 inhibitory activity of VPA was not relevant to the gene expression effects observed in this RN46A assay system. These results suggested inhibition of HDAC1 was most relevant to gene regulation in RN46A cells.

In addition, *HDAC6* was shown to be upregulated by VPA, in RN46A cells, in a prior qPCR study from our laboratory [109] therefore, the effect of HDAC6 inhibition was examined on the VPA-regulated genes. However, our analysis showed tubastatin A (an HDAC6 inhibitor) treatment did not affect any of the selected VPA-regulated genes.

It has been speculated that VPA regulates mood through its non-histone targets and HDACi is responsible for its antitumor properties [110, 111]. However, HDAC inhibitors have been shown to have antimanic effects [112, 113]. VPA, SAHA and MS-275 can normalize mania-like behaviour in the CLOCKΔ19 mouse model [114]. Increased HDAC activity in prefrontal cortex is seen in mania rodent models which can be partially reversed with lithium, VPA and sodium butyrate [115]. VPA DEGs from our study, both HDAC inhibited and non-HDAC regulated, have neuronal functions and are implicated in mood disorders. This suggests that genes altered by VPA are important to mood biology, irrespective of their mechanism of regulation.

## Acknowledgement

This work was supported by the Jim and Mary Carney Charitable Trust, Whangarei, New Zealand, including a Postgraduate Scholarship from this source.

## Conflict of interest

None of the authors report any competing financial interests or other conflicts of interest in relation to this work. The RN46A cell line was a kind gift from Dr Scott Whittemore, Laboratory of Molecular Neurobiology, Louisville, Kentucky, USA.

Supplementary information is available at the Pharmacogenomics Journal’s website.

